# DDA-BERT: leveraging transformer architecture pre-training for data-dependent acquisition mass spectrometry-based proteomics

**DOI:** 10.1101/2024.11.13.623394

**Authors:** A Jun, Pu Liu, Yingying Sun, Jiaying Lin, Xiaofan Zhang, Zongxiang Nie, Yuqi Zhang, Ziyuan Xing, Yi Chen, Tiannan Guo

## Abstract

In data-dependent acquisition mass spectrometry (DDA-MS)-based proteomics, machine learning based rescoring methods are often employed to integrate multiple scores measuring quality of peptide-spectrum matches (PSMs) from different aspects. Existing rescoring tools face limitations incurred by manual feature extraction, shallow machine learning models, and limited training data. Here, we introduce DDA-BERT, a transformer-based end-to-end deep learning model trained with 95 million human spectra for PSM rescoring. DDA-BERT demonstrates superior performance across various sample types, mass spectrometer platforms, trace sample proteomics, and multiple species proteome data. It consistently outperforms state-of-the-art methods of its kind for DDA-MS spectra analysis with up to 103.5% increased protein group identifications. For single cell proteomics, DDA-BERT identifies up to 85.6% more protein groups compared to existing tools. In addition, DDA-BERT can be effectively extended to proteome of non-human species. DDA-BERT offers a robust and scalable solution for enhancing PSM rescoring in proteomics.

## Introduction

In data-dependent acquisition (DDA)-based proteomics, peptide identification typically relies on a protein sequence database search approach. This process begins with the *in silico* digestion of the provided protein sequences to generate theoretical peptide spectra, which are then matched to experimental MS/MS spectra. A machine learning based rescoring approach subsequently integrates multiple scores from database search engines to evaluate the quality of peptide-spectrum match (PSM)^1^. To control false discovery rates (FDR), a target-decoy strategy is commonly used. This method utilizes a decoy database containing reversed or randomized sequences with similar compositional properties to the target database^2^. By sorting and filtering PSM results, the desired FDR is achieved. Final PSM identification results are obtained by setting an FDR threshold.

Over the past few decades, numerous PSM rescoring algorithms have been developed to improve peptide identification reliability. Popular tools include, but are not limited to PeptideProphet^3^, Percolator^4^, Scavenger^5^, and Nokoi^6^. PeptideProphet^3^, a widely used post-processor, applies linear discriminant analysis (LDA) to integrate scores like cross correlation (XCorr) and delta correlation (delta Cn) from database searches into a comprehensive PSM score. Percolator uses a support vector machine (SVM) classifier with semi-supervised learning^4^, distinguishing correct PSMs from incorrect ones by training on decoy and high-scoring target PSMs. Other algorithms, such as gradient boosting decision trees^5^ and L1-regularized logistic regression techniques^6^, have been employed to enhance the classifier’s ability to differentiate between target and decoy PSMs. However, these results rely on shallow machine learning models and manually engineered features that may not sufficiently capture complex nonlinear relationships in peptide-spectrum matching, leading to suboptimal performance. Moreover, these studies train models utilizing sparse training data from a single MS file. Training separate models for individual files prevents them from leveraging patterns learned from other MS files, potentially leading to reduced accuracy and robustness when applied to complex proteomic datasets.

Advancements in deep learning have enabled the development of models that predict peptide characteristics such as MS/MS spectra^7–9^, retention time (RT)^8–10^, collisional cross section (CCS)^8,11^, ion mobility (IM)^8,11^, and binding affinity (BA)^12^. Recently developed rescoring tools, including DeepRescore^13^, InSPIRE^14^, MSBooster^15^, and AlphaPeptDeep^8^, utilize these predictions to generate features. These features, along with other handcrafted features such as spectral correlations, delta RT, normalized spectral angle (SA), and delta CCS, are then fed into rescoring tools such as Percolator to improve the peptide identification in both proteomics and immunopeptidomics. Despite their impressive performance, these tools still rely on handcrafted features and shallow rescoring models, and use deep learning only for generating additional data for feature engineering.

End-to-end deep learning methods offer significant advantages by learning directly from raw data without the need for manual feature engineering, allowing models to capture complex nonlinear relationships more effectively. In *de novo* sequencing tasks, models like DeepNovo^16^, Casanovo^17^, and GraphNovo^18^ have successfully employed end-to-end approaches to determine peptide sequences from mass spectra without relying on handcrafted features, achieving superior performance over traditional methods. This observation naturally raises the question: since end-to-end deep learning methods can achieve better performance on *de novo* sequencing tasks where the sequence space is unconstrained, could they also show improvements in rescoring tasks where the sequence space is constrained? Thus, we introduce DDA-BERT, a pre-trained transformer model that uses an end-to-end approach to learn rescoring functions for peptide-spectrum matches directly from raw MS spectra and peptide sequences, without requiring any additional manual feature engineering.

## Results

### DDA-BERT computational workflow

DDA-BERT adopts a multistage strategy for peptide identification (**Figure 1A**, Methods). Given a DDA-MS file, the database search tool Sage (version 0.14.6) was used to perform a database search. Then a prescoring model, trained on 3706 DDA files containing over 80 million PSMs identified by FragPipe and Sage, was used to score all the PSMs from Sage search results and select qualified PSMs. Next, the rescoring model, which was pre-trained on more than 95 million PSMs selected by the prescoring model—was fine-tuned using the qualified prescored PSMs and then applied to determine the final identification result. The iterative refinement filters out low-confidence candidates generated by the search engine, while pre-training and fine-tuning of the rescoring model improves the reliability and accuracy of the final identification result. Pre-training enables the model to learn from a broader range of PSM samples, enhancing its discrimination capability, while fine-tuning optimizes the model for the specific MS file being analyzed.

**Figure 1.**
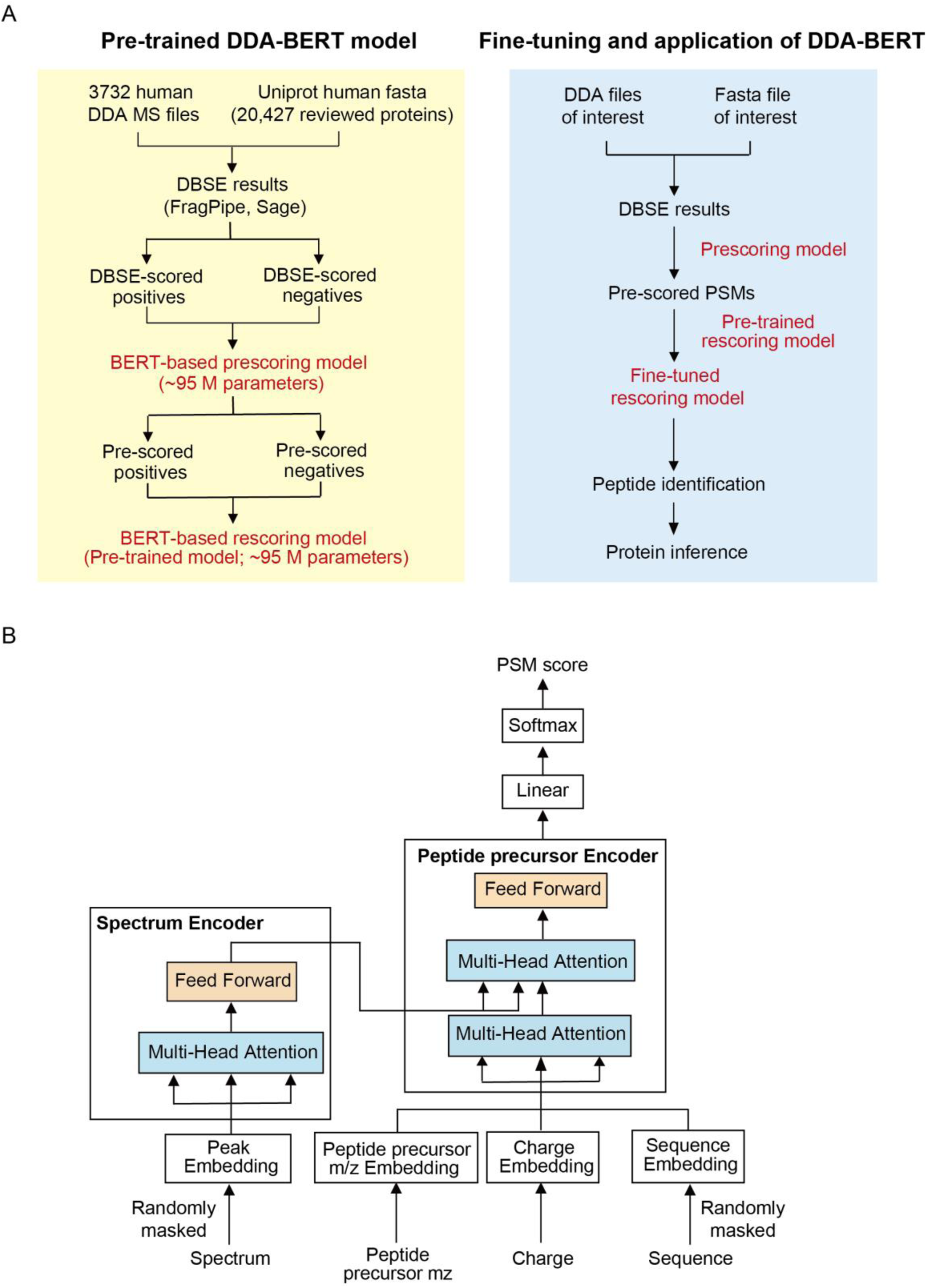
Overview of the pre-training, fine-tuning, and architecture of the DDA-BERT Model. (A) Schematic representation of the pre-training and fine-tuning process of the DDA-BERT model. (B) Detailed architecture of the DDA-BERT model, showing the spectrum encoder and peptide precursor encoder, both incorporating multi-head attention layers for PSM rescoring. DBSE: database search engines.

The rescoring model’s performance on new, unseen data may decline if it memorizes specific sequences rather than understanding the underlying relationship between sequences and spectra. To mitigate this, DDA-BERT employs multi-task learning, introducing an auxiliary task of predicting masked amino acids (**Figure 1B**, Methods). This approach forces the model to grasp the relationship between sequences and spectra, helping it learn more robust and generalizable features. By training the model to both predict the correct PSM and accurately fill in masked amino acids, DDA-BERT achieves good generalization performance.

### DDA-BERT’s rescoring performance across different sample types

The performance of DDA-BERT was compared to two state-of-the-art rescoring tools, namely FragPipe and Sage, at both the PSM and protein levels across two benchmark datasets of different sample types. One dataset comprised human Jurkat T cell samples analyzed on an Orbitrap Exploris 480 mass spectrometer with a 90-minute chromatographic gradient^19^, while the other included colorectal cancer (CRC) FFPE tissue samples analyzed on the same instrument with a 120-minute gradient^20^. The number of PSMs identified by each tool across a range of FDR thresholds (0.1% to 5%) is shown in **Figure 2**. All three rescoring tools identified more PSMs as the FDR threshold increased, with DDA-BERT consistently outperforming the others. At a FDR threshold of less than 0.01, DDA-BERT identified an average of 48,167 PSMs in Jurkat cell, with a 20.9% increase over FragPipe (39,851 PSMs) and a 53.5% increase over Sage (31,371 PSMs) (**Figure 2A**). Moreover, DDA-BERT identified an average of 40,229 peptide precursors (**Figure 2B**), 37,819 peptides (**Figure 2C**), and 6875 protein groups (**Figure 2D**), surpassing FragPipe by 18.8%, 15.1%, and 21.7%, respectively, and outperforming Sage by 47.7%, 42.7%, and 53.9%, respectively. In terms of protein overlaps, over 93.9% of the protein groups identified by FragPipe and more than 98.1% of those identified by Sage were detected by DDA-BERT (**Figure 2E**). In CRC tissue samples, DDA-BERT continued to show superior performance, identifying an average of 90,473 PSMs, which is 73.3% more than FragPipe and 126.6% more than Sage (**Figure 2H**). It also detected significantly more peptide precursors, peptides, and protein groups: 66.0% and 112.4% more peptide precursors (**Figure 2I**), 48.1% and 88.3% more peptides (**Figure 2J**), and 37.7% and 72.2% more protein groups (**Figure 2K**) than FragPipe and Sage, respectively. When compared with MaxQuant^21,22^ and AlphaPept^23^, DDA-BERT achieved a 177.8% increase in PSM identification (**Supplementary Figure 2A**) and a 118.8% increase in protein group identification (**Supplementary Figure 2B**) in CRC tissue samples. Consistent with results from cell line samples, over 92.5% of protein groups identified by FragPipe and more than 97.8% identified by Sage were also detected by DDA-BERT (**Figure 2L**). Additionally, DDA-BERT identified multiple protein groups that were not identified by the other two tools.

**Figure 2.**
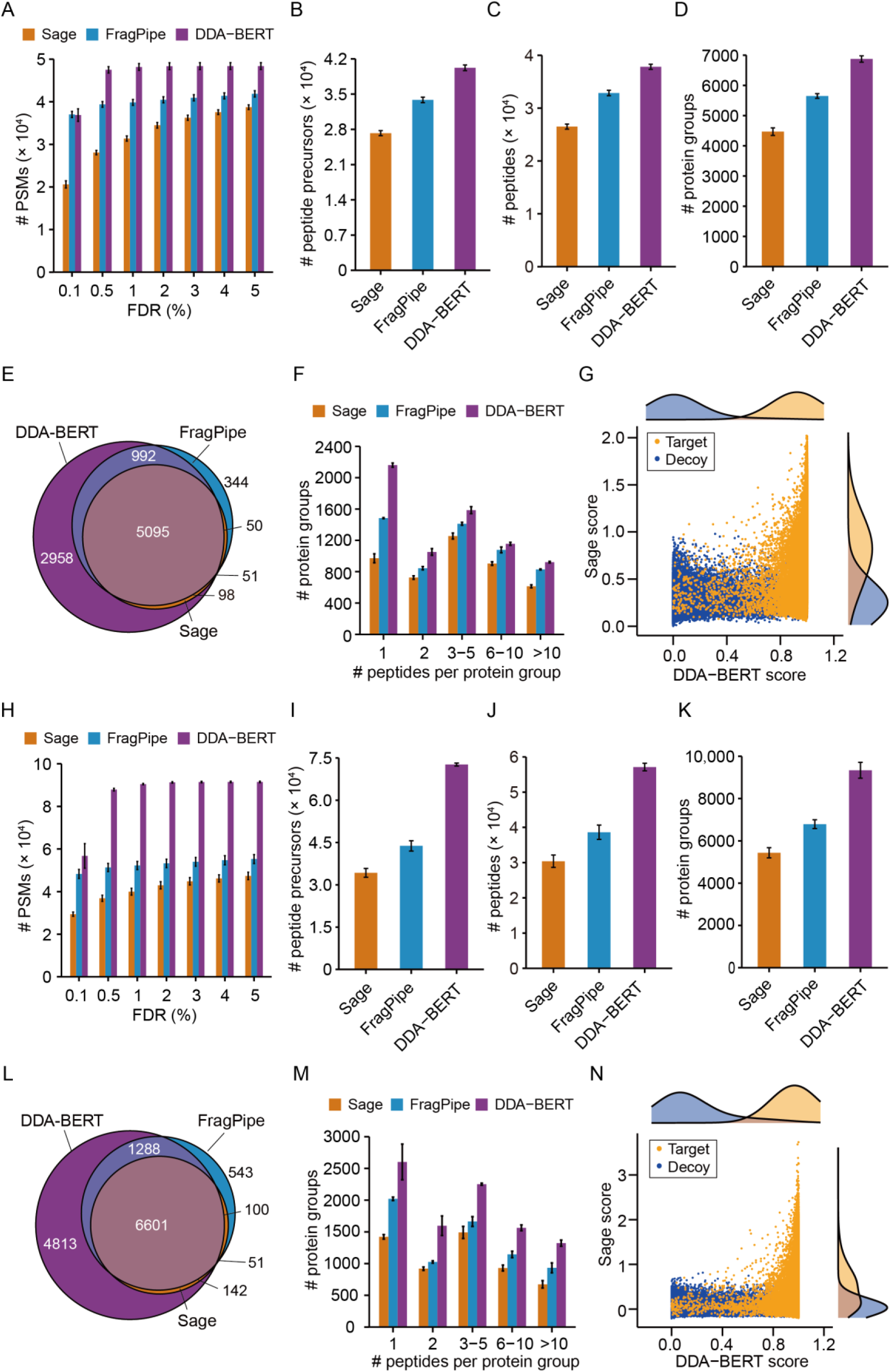
Comparison of the performance of three rescoring tools (DDA-BERT, FragPipe, and Sage) across different sample types. Results for Jurkat cells are shown in panels (A–G), and for colorectal cancer FFPE tissue samples in panels (H–N). (A, H) Bar plots showing the number of PSMs identified at different FDR thresholds. (B, I) Total number of peptide precursors identified by each tool. (C, J) Total number of peptides identified by each tool in the same samples. (D, K) Total number of protein groups identified by these three tools. (E, L) Venn diagrams illustrating the shared and unique protein groups among the three tools. Circle sizes are proportional to the number of protein groups identified by each pipeline. (F, M) Bar plots showing the number of protein groups with 1, 2, 3-5, 6-10, and more than 10 peptides per protein group, identified using Sage (orange), FragPipe (blue), and DDA-BERT (purple). The y-axis represents the number of protein groups identified by each tool; the x-axis indicates the number of peptides per protein group. (G, N) Scatter plots comparing Sage and DDA-BERT scores for target and decoy PSMs in Jurkat cells (G) and colorectal cancer FFPE tissue samples (N), with density plots displayed on the top (Sage) and right (DDA-BERT) axes.

To assess the reliability of protein groups identified by DDA-BERT, we compared the peptide counts per protein group identified by Sage, FragPipe, and DDA-BERT. In Jurkat cells, 69% of the protein groups identified by DDA-BERT were supported by two or more peptides (**Figure 2F**), as were 72% of those in CRC lymph node and tumor deposit samples (**Figure 2M**). Notably, DDA-BERT consistently outperformed Sage and FragPipe in identifying protein groups across both high and low peptide counts. After excluding single-peptide protein groups, DDA-BERT identified 13.2% more protein groups than FragPipe and 34.8% more protein groups than Sage in cell line samples (**Supplementary Figure 3A**). In tissue samples, it identified 41.5% more protein groups than FragPipe and 68.2% more protein groups than Sage (**Supplementary Figure 3B**). Likewise, a comparison of PSM counts (**Supplementary Figures 4A-B**) and peptide precursor numbers (**Supplementary Figures 5A-B**) for each protein group identified by the three tools highlighted the reliability of DDA-BERT in protein group identification. Additionally, we compared the score distributions of target and decoy PSMs from Sage and DDA-BERT. While both exhibited bimodal distributions, DDA-BERT demonstrated better separation between target and decoy PSMs (**Figures 2G and 2N**).

### Rescoring performance on single-cell proteomics

Next, we evaluated the performance of DDA-BERT in proteomics data of trace amount of samples using two published HeLa cell datasets^24,25^. In the first dataset^24^, HeLa cell lysates were analyzed using solid-phase microextraction-assisted capillary zone electrophoresis-mass spectrometry, with injection volumes gradually increasing from 0.1 ng to 10 ng—equivalent to approximately 0.4 to 40 HeLa cells. DDA-BERT identified 523, 1397, 2507, and 9340 PSMs, respectively, which are 2.2%–45.3% more than FragPipe and 5.4%–25.4% more than Sage (**Figure 3A**). In addition, across all injection volume, DDA-BERT also outperformed FragPipe in identifying more peptide precursors (**Figure 3B**), peptides (**Figure 3C**), and protein groups (**Figure 3D**) by 5.6%-41.0%, 5.7%-37.3%, and 16.4%-32.6%, respectively, and surpassed Sage by 5.0%-27.4%, 5.1%-26.3%, and 8.2%-49.1%, respectively. We also tested it in another dataset obtained from a 165-minute chromatographic gradient analysis of a single HeLa cell lysate on an Orbitrap Exploris 480^25^. DDA-BERT identified 14.7% and 35.8% more PSMs (**Figure 3H**), 12.4% and 32.4% more peptide precursors (**Figure 3I**), and 12.3% and 31.8% more peptides (**Figure 3J**) than FragPipe and Sage, respectively. In terms of protein group identification, DDA-BERT outperformed FragPipe by 40.9% and Sage by 85.6% (**Figure 3K**). Similarly, DDA-BERT identified 64.1% and 55.1% more PSMs (**Supplementary Figure 2C**), 148.6% and 136.2% more protein groups (**Supplementary Figure 2D**) than MaxQuant and AlphaPept, respectively. In addition, comparisons of protein overlaps (**Figures 3E and 3L**), multiple-peptide protein group counts (**Supplementary Figure 3C**), peptide distributions per protein group (**Figures 3F and 3M**), PSM and peptide precursor distributions per protein group (**Supplementary Figures 4C and 5C**), and PSM score distributions (**Figures 3G and 3N**) indicated that DDA-BERT outperformed the other tools in peptide identification.

**Figure 3.**
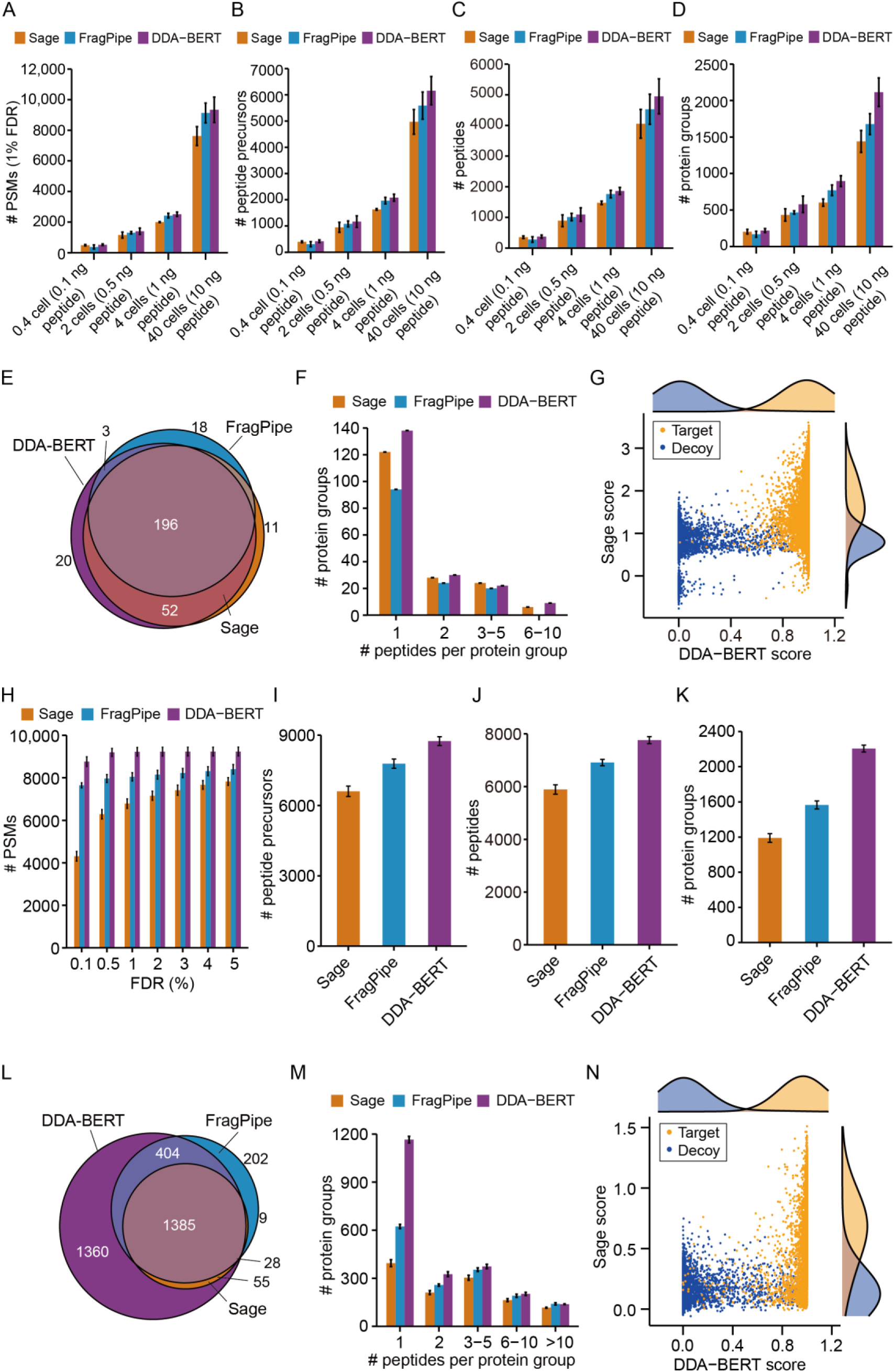
Trace sample proteomics rescoring. Results for 0.4 to 40 HeLa cells are shown in panels (A–G), and for a single HeLa cell are presented in panels (H–N). (A, H) Bar plots showing the number of PSMs identified by DDA-BERT, FragPipe, and Sage. Total number of peptide precursors (B, I) and peptides (C, J) identified by the three tools. (D, K) Total number of protein groups identified by each tool. (E, L) Venn diagrams illustrating the shared and unique protein groups identified in 0.4 HeLa cell and single HeLa cell samples by the three tools. Circle sizes are proportional to the number of protein groups identified by each tool. (F, M) Bar plots showing the distribution of protein groups identified by the three tools in 0.4 HeLa cell and single Hela cell samples, categorized by 1, 2, 3-5, 6-10, and more than 10 peptides per protein group (G, N) Scatter plots comparing target and decoy PSM scores between Sage and DDA-BERT, with density plots displayed for each tool on the top (Sage) and right (DDA-BERT) axes.

### Evaluation in different mass spectrometers and species

We further evaluated DDA-BERT’s performance on different mass spectrometers, including timsTOF Pro and Orbitrap Astral, despite the absence of data from these specific instruments in the training dataset. For DDA data from HCC1806 and HS578T cell lines analyzed with a 60-minute gradient on a timsTOF Pro mass spectrometer^26^, DDA-BERT identified an average of 88,329 PSMs (**Figure 4A**), 52,143 peptide precursors (**Figure 4B**), 38,723 peptides (**Figure 4C**), and 7720 protein groups (**Figure 4D**), outperforming FragPipe by 9.3%, 20.9%, 20.3%, and 31.1%, and Sage by 28.0%, 36.3%, 35.8%, and 43.4%, respectively. For the HeLa cell proteomics data acquired with an Orbitrap Astral mass spectrometer^27^, DDA-BERT identified an average of 75,212 PSMs (**Figure 4H**), 52,851 peptide precursors (**Figure 4I**), 42,394 peptides (**Figure 4J**), and 6759 protein groups (**Figure 4K**), representing a 37.7%, 44.3%, 32.0% and 31.6% increase over FragPipe, respectively; and a 146.4%, 141.0%, 119.3% and 103.5% increase over Sage, respectively. As expected, DDA-BERT also performed well in terms of protein overlaps (**Figures 4E and 4L**), multiple-peptide protein group counts (**Supplementary Figures 3D-E**), distribution of peptides (**Figures 4F and 4M**), PSMs (**Supplementary Figures 4D-E**), and peptide precursors per protein group (**Supplementary Figures 5D-E**), as well as in target/decoy PSM score distribution (**Figures 4G and 4N**). These results highlight DDA-BERT’s robust generalization ability across different mass spectrometers.

**Figure 4.**
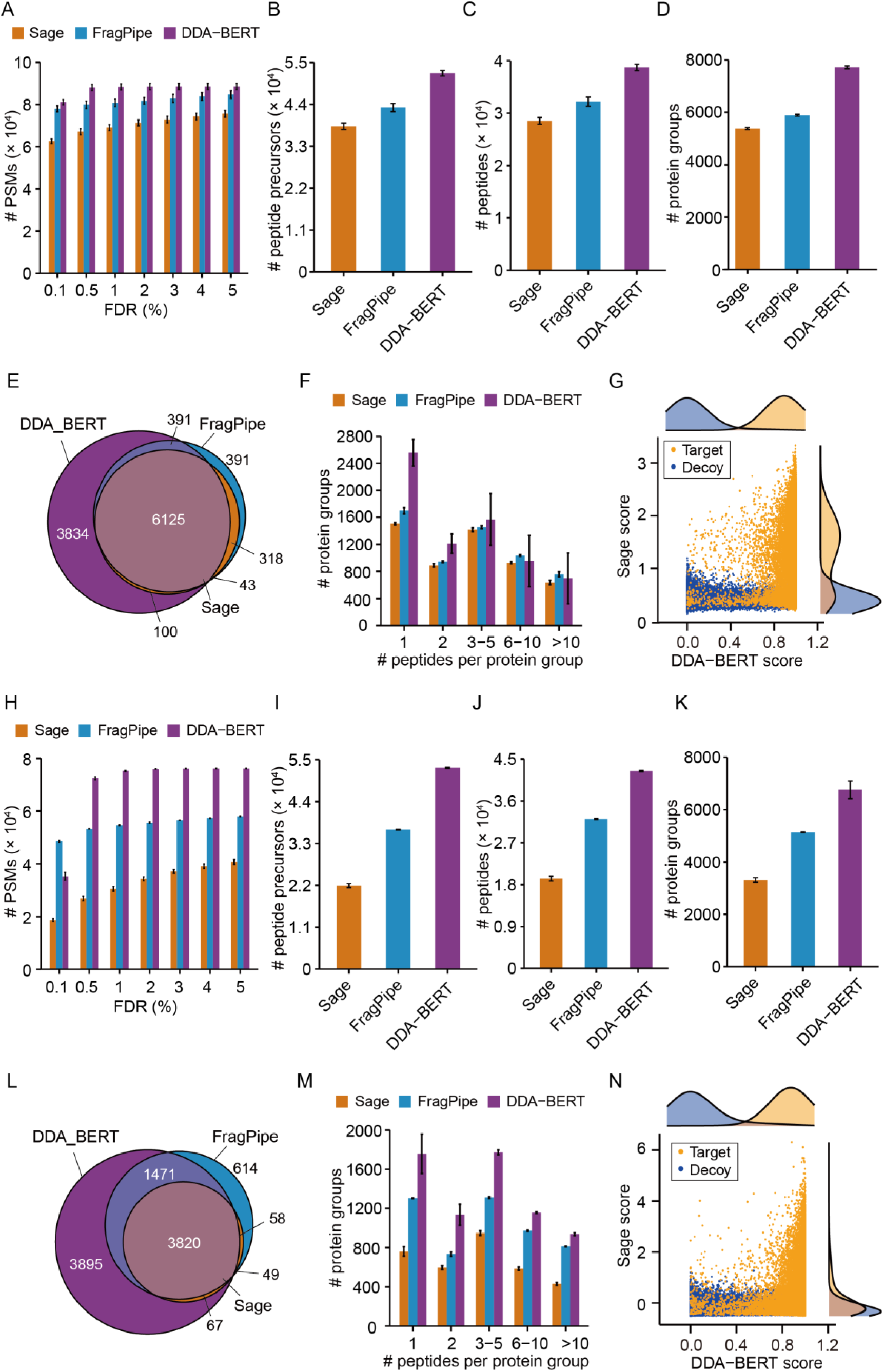
Evaluation of DDA-BERT’s generalization performance across different mass spectrometers. Results for timsTOF Pro mass spectrometer are shown in panels (A–G), and for Orbitrap Astral mass spectrometer are presented in panels (H–N). (A-D) Comparison of the number of PSMs, peptide precursors, peptides, and protein groups identified by DDA-BERT, FragPipe, and Sage using DDA data acquired on the timsTOF Pro mass spectrometer. (H-K) Similar comparisons using data from the Orbitrap Astral mass spectrometer. (E, L) Venn diagrams illustrating shared and unique protein groups identified by these three tools. (F, M) Bar plots displaying the distribution of protein groups with different peptide counts (1, 2, 3–5, 6–10, and more than 10). (G, N) Scatter plots comparing target and decoy scores for Sage and DDA-BERT, with density plots on the top (Sage) and right (DDA-BERT) axes.

Since the DDA-BERT model was trained on human MS data, we further assessed its generalization performance with a mouse dataset derived from liver samples of 12-month-old mice, acquired on a Q Exactive HF mass spectrometer over a 180-minute run^28^, where it identified an average of 68,213 PSMs (**Figure 5A**), 39,964 peptide precursors (**Figure 5B**), 27,037 peptides (**Figure 5C**), and 3453 protein groups (**Figure 5D**), exceeding FragPipe by 8.5%, 26.8%, 26.3%, and 3.8%, and surpassing Sage by 13.7%, 30.2%, 31.6%, and 12.2%, respectively. Anlysis of other metrics, including protein overlaps (**Figure 5E**), multiple-peptide protein group counts (**Supplementary Figure 3F**), distributions of peptide (**Figure 5F**), PSM (**Supplementary Figure 4F**), and peptide precursor per protein group (**Supplementary Figure 5F**), as well as PSM score distributions (**Figure 5G**), aligned with findings from HeLa cell proteomics data obtained using the Orbitrap Astral, further supporting the strong generalization ability of DDA-BERT.

**Figure 5.**
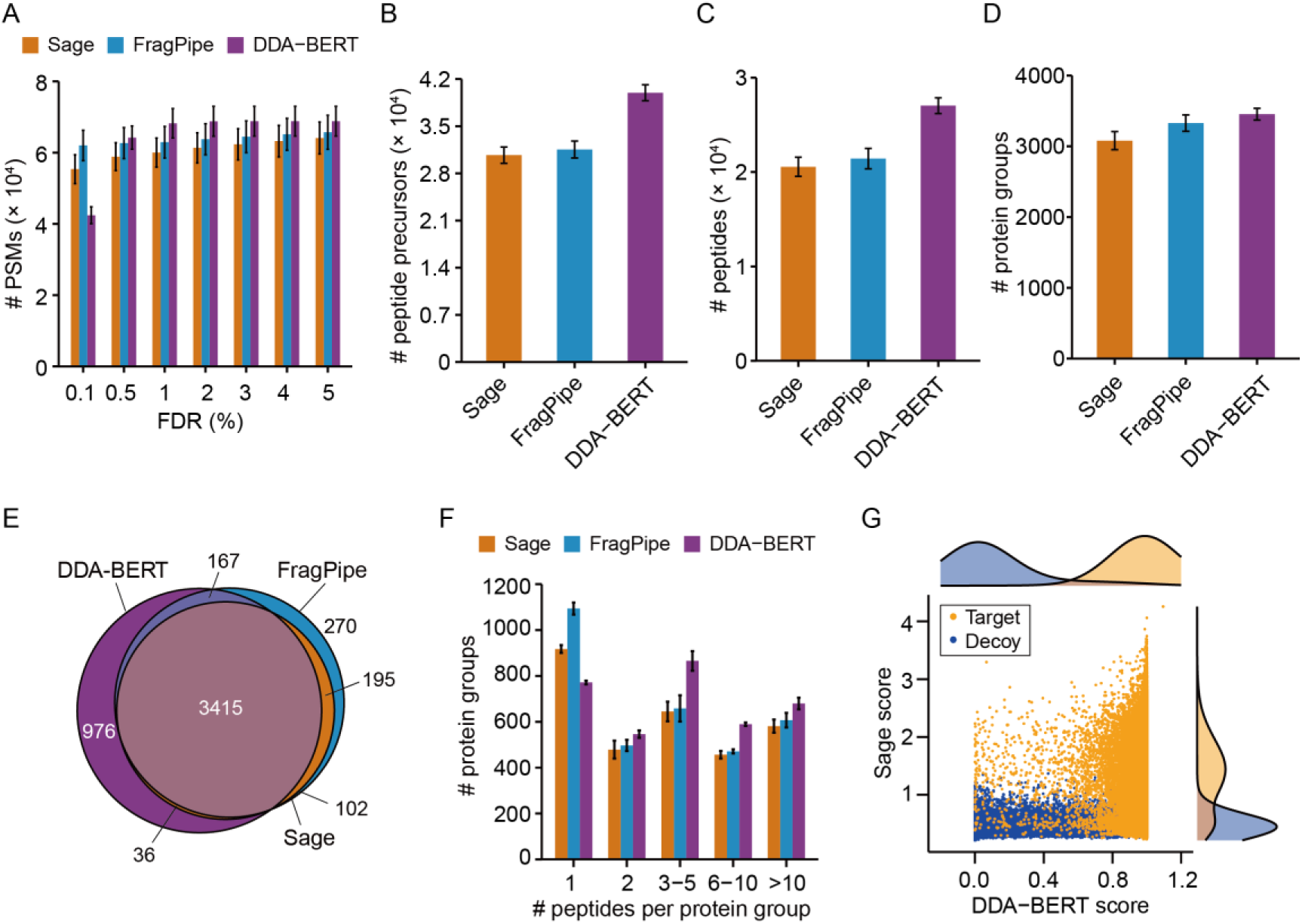
Evaluation of DDA-BERT’s generalization performance on a mouse dataset. (A-D) DDA-BERT’s performance in identifying PSMs (A), peptide precursors (B), peptides (C), and protein groups (D), as well as its protein overlap (E) with FragPipe and Sage. (F) Bar plots showing the number of protein groups identified with 1, 2, 3-5, 6-10, and more than 10 peptides. (G) Scatter plot comparing target and decoy PSM scores between Sage and DDA-BERT, with density plots displayed along the top (Sage) and right (DDA-BERT) axes.

## Discussion

Inspired by the success of the Transformer model in *de novo* peptide sequencing^17,29,30^, we developed a pre-trained Transformer-based model specifically designed for PSM rescoring to enhance peptide identification in DDA data. Trained on over 95 million spectra, DDA-BERT effectively captures complex nonlinear relationships in peptide-spectrum matching, yielding superior discrimination capabilities. Fine-tuning further enables the model to adapt to the unique characteristics of specific mass spectrometry files. Additionally, DDA-BERT is trained directly on raw data, eliminating the need for manual feature extraction.

DDA-BERT consistently outperforms other tools across various sample types and mass spectrometry platforms—from trace samples to mouse proteomics, with a user-friendly interface to facilitate its adoption by researchers. However, it does have certain limitations, such as the requirement for at least one graphics processing unit (GPU) to run, which can be a barrier for users with limited computing resources. This is because the BERT models, utilizing more signals from the raw MS data and relying on much more parameters, require much more computing than other available software tools. Future improvements focus on evaluating DDA-BERT’s performance across various application scenarios, including HLA immunopeptidomics and neoantigen discovery, expanding its dataset to cover additional species and high-quality spectra. Planned extensions also include applying DDA-BERT to small molecules and lipid analysis, as well as quantitative analysis in labeled proteomics. This study highlights the significant potential of pre-trained DDA-BERT model in DDA-MS data analysis. DDA-BERT is poised to significantly enhance our understanding of complex biological systems, aid in the discovery of new clinical therapeutic targets, and advance the exploration of understudied proteins and peptides.

### Online methods

#### Dataset preparation

Database searches were performed on 3732 human MS files from 29 datasets using Sage v0.14.6^31^ and FragPipe v21.1^32^. Raw MS/MS files were first converted to mzML format using msconvert GUI (ProteoWizard, v3.0.19014). Searches were then conducted against a protein database with 20,427 SwissProt human protein sequences (ver. 2023/10), 96 common contaminants and their pseudo-reversed sequences. Parameters included peptide lengths of 7–50, 20 ppm mass tolerances, trypsin digestion with a maximum of two missed cleavages, variable modifications (oxidation of methionine, N-terminal acetylation, deamidation of asparagine and glutamine), and fixed modification of carbamidomethylated cysteine. The FDR for both PSM and protein were set at 0.01. Detailed parameters are listed in **Supplementary Table 1**.

#### Training of prescoring model

PSMs were identified at a 1% FDR using FragPipe and Sage on 3732 mass spectrometry files (termed “identification result”). Sage also provided the top 10 candidate PSMs and scores per scan (termed “Sage search result”). The target PSMs from the intersection set of the identification result and Sage search result were used as positive training instances. Decoy PSMs from Sage search results were used as native training instances, with low-score decoy PSMs randomly removed to achieve a 1:1 ratio with positives. From over 80 million extracted PSMs, the prescoring model was trained and validated using an independent validation dataset.

#### Pre-training of rescoring model

Each PSM in the Sage search results was scored by the prescoring model. Target PSMs with scores above 0.5 were designated as positive instances, while decoy PSMs were marked as negative instances, with low-score decoy PSMs randomly removed to achieve a 1:1 ratio with positive instances. The rescoring model was pre-trained on over 95 million extracted PSMs and validated using an independent validation dataset.

#### Fine tuning of rescoring model and Identification

At the peptide identification stage, Sage was first used to perform a database search on the raw mass spectrometry file and generate up to 100 peptide sequence candidates per fragmentation spectrum. The prescoring model then scored all PSMs from the Sage search results. Target PSMs with scores greater than 0.5 were chosen as positive instances, while an equal number of high-scoring decoy PSMs were used as negative instances. The pre-trained rescoring model was fine-tuned on this dataset. The fine-tuned model re-scored PSMs and calculated 1% FDR. Protein inference was performed using the method described by Mann et al. to obtain the final protein identification results^23^.

#### Model Architecture

DDA-BERT comprises two interconnected transformer encoders: a spectra encoder for processing peak m/z and intensity data from MS2 scans, and a peptide precursor encoder for processing peptide sequences along with peptide precursor m/z and charge (**Figure 1B**). Similar to the approach used in Casanovo^17^, DDA-BERT uses sinusoidal position embeddings and learned linear layers to encode MS2 m/z values and intensities, while also using learned embedding layers for the peptide precursor’s m/z and charge. The sum of the learned embedding of each amino acid and its position embedding is concatenated to encode the amino acid sequence. Causal masking, however, is not used in peptide precursor encoder since DDA-BERT is not designed for autoregressive tasks. Cross-attention is applied between two encoders to allow the model to combine the spectral data with the peptide precursor information in a context-aware manner, and align and correlate the relevant features from both encoders.

To enhance the model’s ability to understand the complex relationships between the MS/MS spectrum and the peptide sequence rather than memorize frequently occurring peptide sequences in the training set, a masked amino acid prediction loss function is employed. The model is configured with nine layers, an embedding size of 768, and sixteen attention heads, totaling approximately 95 million parameters. Training was conducted with a batch size of 128 spectra, a weight decay of 5 × 10⁻⁵, and a peak learning rate of 5 × 10⁻⁵. The learning rate followed a linear increase over 150k warm-up steps, followed by cosine decay. Training was performed on 16 NVIDIA-A100 GPUs over 30 epochs, taking approximately two days. The model weights selected for testing corresponded to the epoch with the lowest validation loss. Further details on training and hyperparameters can be found in **Supplementary Table 2**.

#### Metrics and benchmarks

We evaluated DDA-BERT against state-of-the-art tools like FragPipe and Sage, using seven publicly available datasets, ensuring consistent FDR estimation across all pipelines. Additionally, the same protein inference method was employed to facilitate comparability in protein identification results. Details of the benchmark datasets are as follows:

Dataset 1: Lysates from Jurkat cells were analyzed using an Orbitrap Exploris 480 mass spectrometer with a 90-minute chromatographic gradient. The raw MS/MS files are available on PRIDE (accession code: PXD036024)^19^.

Dataset 2: Formalin-fixed, paraffin-embedded (FFPE) tissue samples from colorectal cancer patients were analyzed using an Orbitrap Exploris 480 mass spectrometer with a 120-minute gradient. The dataset is accessible via PRIDE (accession code: PXD041625)^20^.

Dataset 3: Proteomics data for samples containing 0.4, 2, 4, and 40 cells were acquired using a Q Exactive HF mass spectrometer. This dataset can be accessed through PRIDE (accession code: PXD047627)^24^.

Dataset 4: Single HeLa cell lysate analyzed using an Orbitrap Exploris 480 mass spectrometer. The dataset can be accessed on PRIDE (accession code: PXD041879)^25^.

Datasets 5-6: DDA data from HCC1806 and HS578T cell lines were generated using a timsTOF Pro mass spectrometer with a 60-minute chromatographic gradient. Additionally, HeLa cell digests (200 ng) were analyzed using an Orbitrap Astral mass spectrometer. These datasets are available on PRIDE under accession codes PXD041391^26^ and PXD046453^27^, respectively.

Dataset 7: Liver tissue samples form mice were analyzed on a Q Exactive HF mass spectrometer using a 180-minute chromatographic gradient. This dataset is available via PRIDE (accession code: PXD035255)^28^.

## Data availability

Proteomic datasets used in this study are accessible from the ProteomeXchange Consortium and the PRIDE Partner Repository^33^ or from the iProx Repository^34,35^, identified by the following access codes: PXD023209, PXD019643, PXD031389, PXD029202, PXD031323, PXD021371, PXD014199, PXD008463, PXD021089, PXD025626, PXD040391, PXD021169, PXD041066, PXD010966, PXD020612, PXD020630, PXD028680, PXD018346, PXD023634, PXD030569, PXD026635, PXD028785, PXD021164, PXD031311, PXD020659, PXD023533, PXD040732, IPX0001400000, and IPX0005714000. Additionally, the access codes for the benchmark dataset used to assess the performance of the DDA-BERT model are PXD036024, PXD046058, PXD041625, PXD047627, PXD046453, PXD041391, and PXD035255.

## Software development and availability

DDA-BERT code is available freely and as open source at https://github.com/guomics-lab/DDA-BERT. Executable versions can be accessed via https://dda-bert.guomics.com/. DDA-BERT is compatible with Windows desktop computers and also supports the Linux operating system. The current processing time for a 90-minute gradient DDA file from the Orbitrap Exploris 480 is approximately 1.3 h using one NVIDIA A100 GPU and 10 CPUs.

## Acknowledgements

This work is supported by grants from National Key Research and Development Project Subject (Grant No. 2021YFA1301603), “Pioneer” and “Leading Goose” R&D Program of Zhejiang (2024SSYS0035), the Zhejiang Province Leading Geese Plan (2024SSYS0035), and the Key Research and Development Program of Zhejiang Province (Grant No. 2022C03037). We gratefully acknowledge the support of Westlake University High-Performance Computer Center. We thank the Research Center for Industries of the Future (RCIF) at Westlake University for partially supporting this work. Declaration of generative AI and AI-assisted technologies in the writing process during the preparation of this work the authors used ChatGPT to improve language and readability. After using this tool/service, the authors reviewed and edited the content as needed and took full responsibility for the content of the publication.

## Author contributions

T.G. acquired funding and supervised the project. T.G. and Y.C. conceptualized the research and provided oversight. Y.C. designed the methodology and laid the foundation for research implementation. J.A, P.L., and J.L. managed the overall experimental pipeline. Z.N. developed the data preprocessing component and the DDA-BERT software system. J.A, J.L., X.Z., Y.Z., Y.S., and Z.X. curated the data. J.A, P.L., J.L., and X.Z. conducted model parameter tuning and performed analyses. J.A designed the benchmark experiments and drafted the initial manuscript with contributions from J.L., P.L., and X.Z., J.A, Y.C., Y.S., and P.L. revised the manuscript. All authors have reviewed and approved the final version of the manuscript.

## SUPPLEMENTAL INFORMATION

**Supplementary Figure 1.**
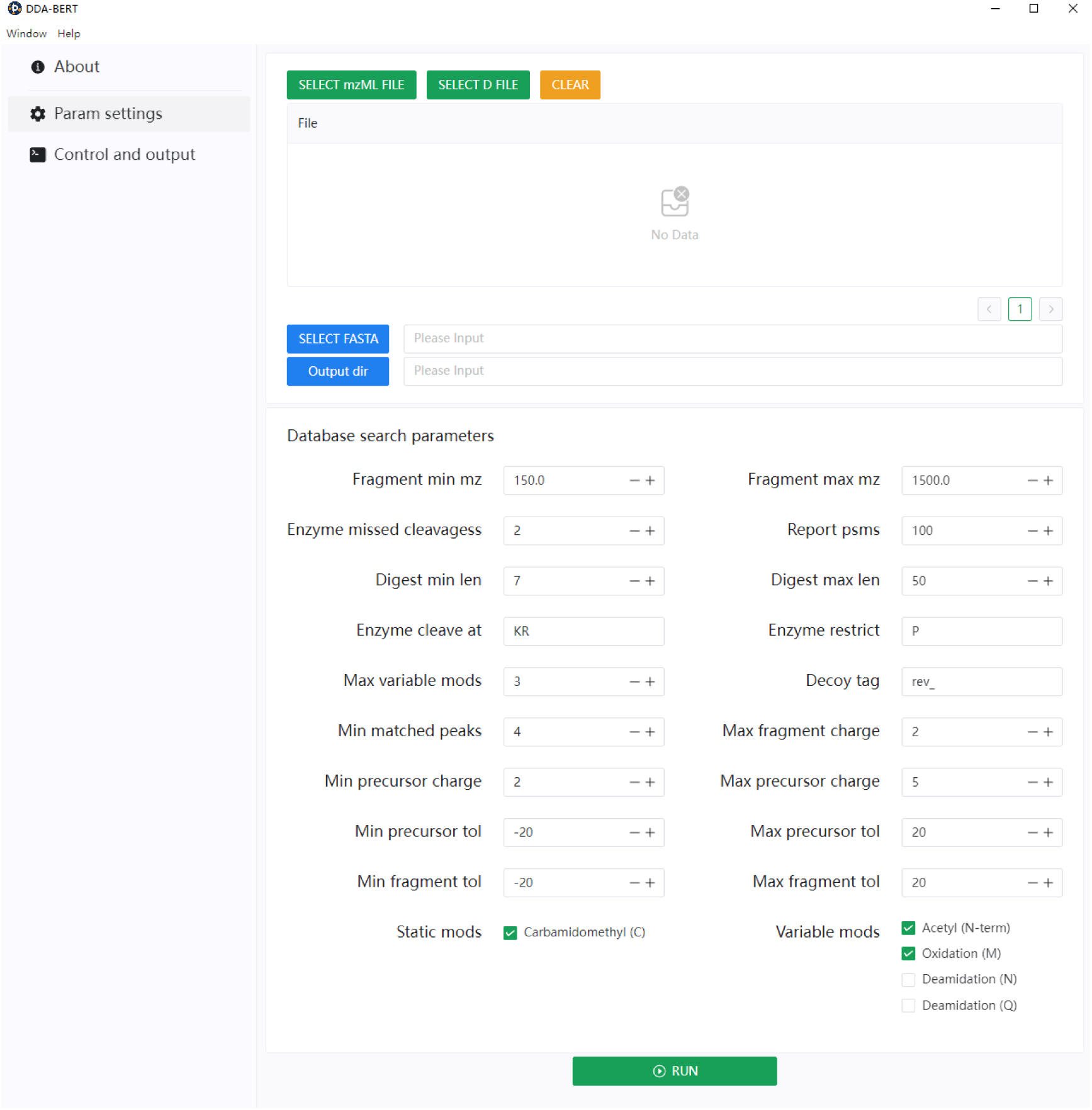
Graphical user interface of DDA-BERT. The interface is divided into three main sections: Input, Param Settings, as well as Control and Output. In the Input section, users select the FASTA file and specify the output directory paths. The Param Settings section allows users to configure detailed database search parameters, including fragment mass range (min and max), number of missed enzyme cleavages, peptide length range, enzyme cleavage specificity, maximum variable modifications, and precursor/fragment charge limits. Users can also define static and variable modifications, as well as tolerance levels for precursor and fragment ions. The Control and Output section provides options to execute the run by pressing the “RUN” button and monitor progress.

**Supplementary Figure S2.**
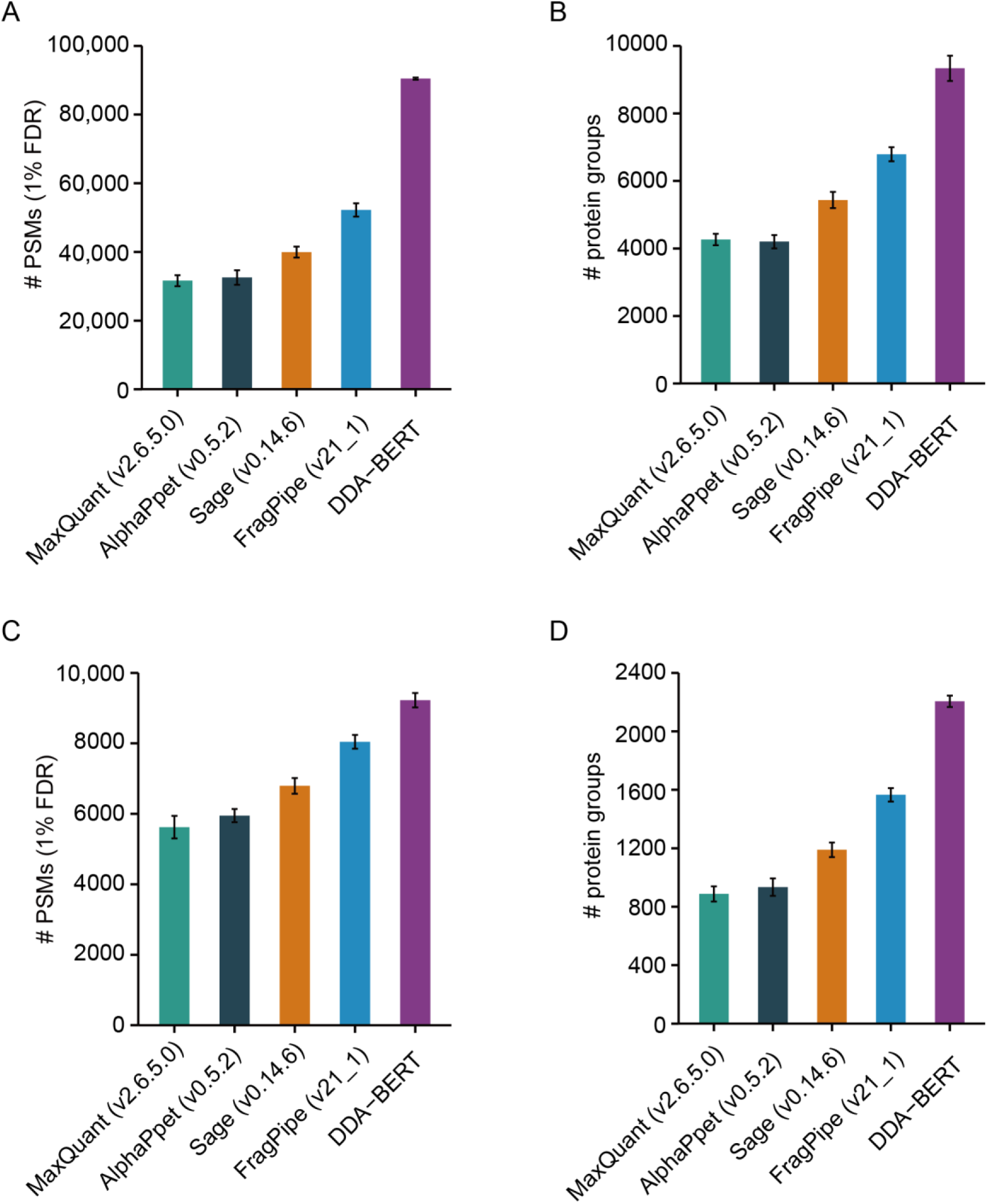
Comparison of PSMs and protein groups identified by MaxQuant, AlphaPept, Sage, FragPipe, and DDA-BERT. (A, C) PSMs identified at 1% FDR in colorectal cancer FFPE tissue (A) and single Hela cell (C). (B, D) Protein groups identified by each tool in the same samples.

**Supplementary Figure S3.**
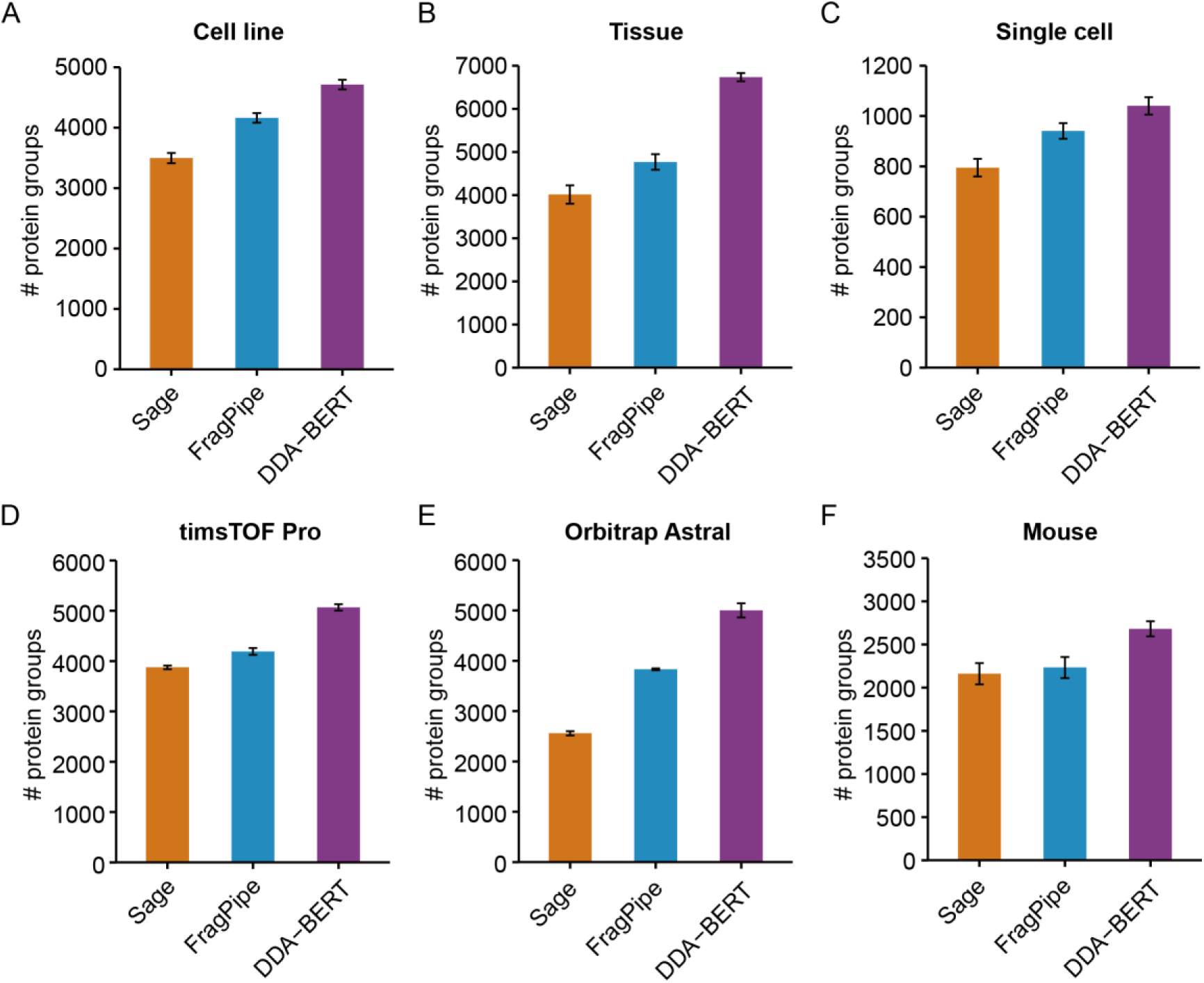
Comparison of the number of protein groups with more than one peptide identified by Sage, FragPipe, and DDA-BERT across various sample types and mass spectrometers. Bar graphs (A-C) show the number of protein groups identified by Sage (orange), FragPipe (blue), and DDA-BERT (purple) in Jurkat cell (A), colorectal cancer FFPE tissue samples (B), and single HeLa cell (C). Graphs (D-E) exhibit DDA-BERT’s performance on the timsTOF Pro (D) and Orbitrap Astral (E) mass spectrometers. Graph (F) displays the protein groups identified using the three rescoring tools in a mouse dataset. The y-axis represents the number of protein groups identified with more than one peptide.

**Supplementary Figure S4.**
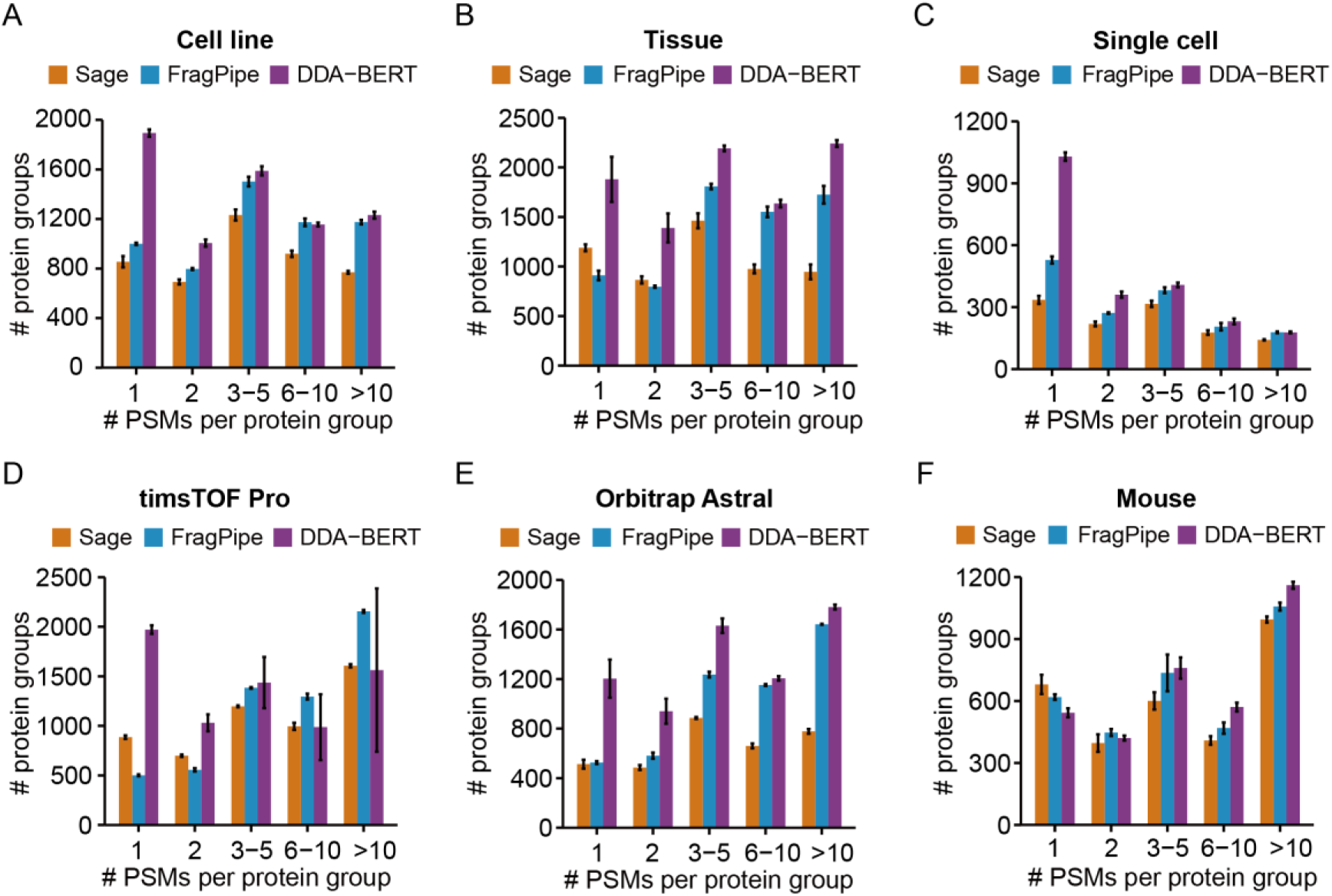
Comparison of PSM distributions among protein groups identified by Sage, FragPipe, and DDA-BERT. Bar plots (A-F) display the distribution of protein groups identified with 1, 2, 3-5, 6-10, and more than 10 PSMs across different datasets or experimental conditions for Sage (orange), FragPipe (blue), and DDA-BERT (purple). Panels correspond to: Jurkat cells (A), colorectal cancer FFPE tissue samples (B), single Hela cell (C), timsTOF Pro mass spectrometer (D), Orbitrap Astral mass spectrometer (E), and a mouse dataset (F). The y-axis represents the number of protein groups identified by each tool, and the x-axis shows the distribution of PSMs per protein group.

**Supplementary Figure S5.**
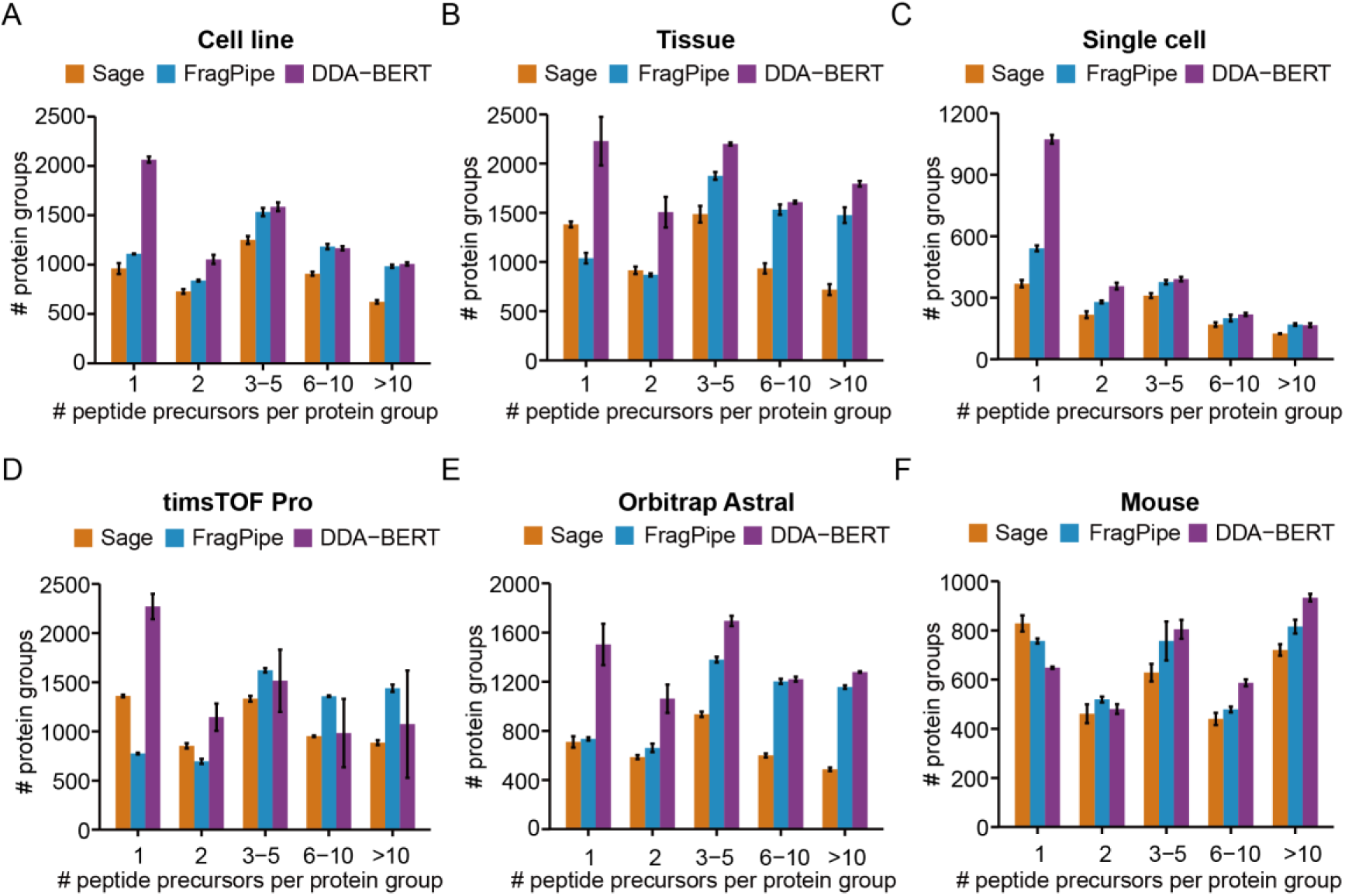
Comparison of peptide precursor distributions among protein groups identified by Sage, FragPipe, and DDA-BERT. Bar plots (A-F) display the distribution of protein groups identified with 1, 2, 3-5, 6-10, and more than 10 peptide precursors across different datasets or experimental conditions for Sage (orange), FragPipe (blue), and DDA-BERT (purple). Panels correspond to: Jurkat cells (A), colorectal cancer FFPE tissue samples (B), single Hela cell (C), timsTOF Pro mass spectrometer (D), Orbitrap Astral mass spectrometer (E), and a mouse dataset (F). The y-axis represents the number of protein groups identified by the three tools, and the x-axis shows the distribution of peptide precursors per protein group.

